# TU_MyCo-Vision: A Deep Learning Tool for Detection of Cell Morphologies in Fungal Microscopic Images

**DOI:** 10.1101/2025.07.23.666349

**Authors:** Kartik J. Deopujari, Matthias Schmal, Caroline Danner, Zainab Abdul Qayyum, Jordy T. Zwerus, Julian Kopp, Mihail Besleaga, Roghayeh Shirvani, Astrid R. Mach-Aigner, Robert L. Mach, Christian Zimmermann

## Abstract

Morphological switching in response to environmental stimuli is a well-known phenomenon in fungi, leading to diverse morphotypes. Microscopic observation remains a widely used approach to study these phenotypes. However, variation in sample preparation and operators skill can limit the scale of sample processing or introduce bias. Although several image-based cell detection tools have been developed, most are tailored to specific applications or limited to a particular taxon. To address the need for a tool applicable to the polymorphic, yeast-like fungus *Aureobasidium pullulans,* and with potential applicability to other taxa, we developed TU_MyCo-Vision, an Ultralytics YOLO (You Only Look Once) based object detection tool for identifying 13 fungal morphotypes in bright-field microscopic images.

The tool integrates a YOLOv11m-based object detector trained on a custom dataset of 1,504 annotated images and a standalone graphical user interface that enables downstream data analysis and visualization of results. The best-performing model (Zulu_s3) achieved a mean precision of 73.4%, a recall of 66.5%, a mean average precision at 50% IoU (mAP@50) of 73.5%, and a mean average precision at varying IoU thresholds between 50 to 90% IoU (mAP@50-95) of 54.5% across all 13 classes. The single-group analysis pipeline was validated on a 90-image test set, generating six quantitative summaries, including absolute counts, relative and mean relative abundance plots, stacked bar plots, and clustered heatmaps. Multi-group evaluation on previously unseen datasets comprising *Candida albicans*, *Komagataella phaffii*, and *Aspergillus niger* spores demonstrated the tool’s potential applicability to other genera.

TU_MyCo-Vision is distributed as a fully packaged, cross-platform executable, eliminating the need for environment setup or manual installation of dependencies. Built entirely on open-source frameworks, it provides a foundational and potentially extensible solution for automated fungal morphology detection and analysis.

**Author Summary:** We developed TU_MyCo-Vision to address challenges in fungal microscopic imaging. Fungi, such as *Aureobasidium pullulans*, display a remarkable ability to switch cell shapes (up to thirteen in this species alone) depending on their environment. While microscopy remains a popular method for observing these changes, manual analysis is limited by individual expertise and the number of images that can be processed, often making results subjective and difficult to scale. To overcome these challenges, we built an Ultralytics YOLOv11-based cell detector that can automatically detect and categorize thirteen fungal cell shapes from brightfield microscopic images. We designed TU_MyCo-Vision to be accessible, with a simple graphical user interface, integrated data analysis suite, and distribution as a standalone application for both Windows and macOS, so it can be used even by those with limited computational skills. Our tool demonstrated strong performance, achieving over 73% precision. Importantly, it also worked well on images from other fungal species, showing potential to be further developed as a general fungal cell morphology tool. We hope TU_MyCo-Vision will contribute to making standardized, high-throughput phenotyping of fungi accessible to a broader community.

## Introduction

The kingdom of fungi is well known for its diversity, especially when it comes to mode of nutrition, habitat, and morphology. Although diversity introduces its challenges in studying fungi, the identification of morphologies through direct microscopic observation remains a commonly used method (1). An interesting phenomenon in some fungi is their ability to exhibit different morphologies throughout their life cycle. These changes in morphology may occur in response to environmental stimuli or play a role in pathogenicity (2). A good example is the morphological switching in *Candida albicans,* where generally the hyphal and pseudohyphal morphologies are considered pathogenic over yeasts, which are regarded as commensal (2,3). A Similar tendency to switch morphology is also observed in soil-borne pathogenic fungi *Coccidioides immitis* and *Coccidioides posadasii,* in which spherules form through the progressive enlargement of arthroconidia that enter the body via inhalation (4). This highlights the importance of morphological assessment, particularly in clinical settings, where treatment decisions may depend on the presence of septate hyphae in *Aspergillus* versus non-septate hyphae in *Mucorales*, as observed through micromorphological analysis. While the microscopic examination has its own advantages, such as its wide applicability and rapid results, it is also sensitive to factors like the quality of sample preparation and operator skill (5–7).

To address conventional limitations, several deep learning tools have been developed for fungal detection in microscopy. For example, Koo et al. introduced a YOLOv4-based object detector that automates the KOH (potassium hydroxide) examination for rapid hyphae detection in microscopic images (6). Candescence employs a Fully Convolutional One Stage (FCOS) architecture to identify and classify nine *C. albicans* morphologies from DIC (Differential Interference Contrast) microscopic images (8). For edible fungi, CCHA-YOLO enables automated and efficient detection of mycelium clamp connections and hyphae autolysis, processes that are typically time-consuming but essential for examining mycelial growth and aging. CCHA-YOLO achieved a high mAP50-95 of 89.02% and includes a web interface to improve accessibility (9,10).

Separately, an improved YOLOv5-based tool was developed to detect downy mildew spores in natural scenes, facilitating early outbreak prediction through automated spore detection (11).

Fungal polymorphism is important not only in clinical settings but also in biotechnological applications. *Aureobasidium pullulans* is a polyextremotolerant black yeast-like fungus with significant biotechnological importance. It is known to survive across a wide spectrum of extreme environments, including hypersaline, highly acidic or basic, cold, and oligotrophic conditions. *A. pullulans* is well known for producing pullulan, along with other metabolites such as heavy oils and polymalic acid. Members of the *Aureobasidium* genus exhibit remarkable phenotypic plasticity in response to environmental conditions (12).

This leads to the development of various morphotypes, which include yeast-like cells, blastoconidia, chlamydospores, hyphae, pseudohyphae, swollen cells, and septate cells (13–15). Li et al. described several morphological forms in *A. pullulans* NG, including yeast-like blastospores (YLB), yeast-like cells (YL), swollen cells (SC), septate swollen cells (SSC), chlamydospore-like cells (CH), hyphal forms (HY), and meristematic structures (MS) (16). Among these, the SC form plays a key role, acting as a relatively stable intermediate that arises from YL cells. This form is notable for producing key metabolites such as unpigmented pullulan and subsequently differentiating into melanin-rich forms like CH, HY, and MS as the organism progresses into later growth stages. A practical application of this was demonstrated by (15), where they demonstrated the role of citric acid in controlling the cellular differentiation in *A. pullulans* NG, which led to efficient pullulan biosynthesis due to the growth in the cells in swollen cell form.

In the course of our own cultivation experiments with *A. pullulans,* we observed a high degree of morphological variability across conditions and strains and generated a large volume of brightfield microscopy images. While several fungal cell detection tools are available, most are limited to specific taxa or morphologies and lack accessibility for users without computational expertise, preventing their application for our purposes. Thus, we developed TU_MyCo-Vision, an object detection tool based on the Ultralytics YOLOv11m architecture, aiming at detecting thirteen morphotypes in *Aureobasidium*.

The pretrained YOLOv11m model was re-trained using a curated dataset of 1504 brightfield microscopic images, primarily representing *A. pullulans*, with additional images from *Aureobasidium melanogenum* and *Trichoderma reesei* to improve generalizability. A hybrid annotation approach combining expert labeling and model-assisted auto-labeling was used for dataset curation. A 5-fold cross-validation strategy was used for training and evaluation. The dataset was iteratively improved during the study, leading to improved performance of the final model. The final model was tested on images of *Komagataella phaffii*, *Aspergillus niger*, and *C. albicans*, demonstrating potential applicability to genera not part of the training dataset. To improve accessibility, we integrated the model into a graphical user interface (GUI) along with a custom data analysis pipeline, enabling users to interpret results without requiring programming skills. The GUI tool enables fungal cell detection, visualization, and downstream analysis, including single and multi-group comparisons, class exclusion for focused morphology-specific analysis, and interactive plots for easier visualization.

## Materials and Methods

### A. Dataset generation

A microscopic image dataset of *A. pullulans* EXF-150, 3374, 3519, 3750, 4010, 5628, 6298, 6519, 8127, 8128, and 10632 (17) and NBB 7.2.1 (18) was generated through a combination of different cultivation experiments aimed at studying the morphology of *A. pullulans*. To increase dataset diversity and improve model generalizability, microscopic image data of *T. reesei* and *A. melanogenum* were included.

#### A.1 Cultivation of *A. pullulans*

*A. pullulans* strains were cultivated in *A. flavus* and *A. parasiticus* medium (AFP), Archimycetes medium (ARCH), Brain Heart Infusion medium (BHI), boiled rice medium, potato dextrose medium (PD), yeast peptone dextrose medium (YPD), M17 medium, malt extract medium (MEX), Muller Hinton broth (MH), and minimal medium. Refer to “S1 Table 1” for detailed medium composition. 100 ml Flasks containing 25 ml of respective cultivation medium and strain were incubated for different time periods, and incubation temperatures ranging from 14 °C to 34 °C, and at different shaking speeds ranging from 200 – 220 rpm. Media compositions adapted from (19)

##### A.1.1 Cultivation of *A. melanogenum*

*A. melanogenum* ATCC 42023 was incubated on MEX plates at 22 °C for 5 days. Yeast-like growth was scraped off with a sterile inoculation loop and resuspended in 0.8% NaCl. 25 ml of fresh medium was inoculated with the suspension with a starting OD600 of 0.05 and incubated at 24°C at 200 rpm for a period between 24h and 168h, approximately.

##### A.1.2 Cultivation of *T. reesei*

*T. reesei* strains RL-P37 (NRRL 15709) and Rut-C30 were cultivated on PD agar plates at 30°C until sporulation. The conidia were harvested and resuspended in 0.8 %(w/v) NaCl, 0.05 % (v/v) Tween-80. 6-well culture plates, with each well containing 5 ml of growth medium, were inoculated to a density of 10^9^ conidia/L. The cultures were incubated at 30 °C for 20 hours under static conditions. Three types of media were used for cultivation: Mandels–Andreotti (MA) medium supplemented with either 1% (w/v) lactose or 1% (w/v) glucose as the sole carbon source, and MEX medium.

##### A.1.3 Cultivation of *K. phaffii GS115*

The pre-culture medium (100 mL/L of sterilized 0.1 M potassium phosphate buffer pH 6.0, 13.4 g/L yeast nitrogen base without amino acids and with ammonium sulfate, 5 g/ L (NH4)2SO4, 400 mg/L biotin, 20 g/L glycerol and 100 µg/ml Zeocin) was inoculated with a fresh cryo stock, and incubated at 30 °C, 230 rpm for 24 h. Next, the pre-culture was used to inoculate the batch medium (10% of the batch media volume). All cultivations were performed in a Minifors 2 bioreactor system (max. working volume: 2 L; Infors HT, Basel, Switzerland). Process control and feeding were performed using EVE software (Infors HT, Bottmingen, Switzerland). The batch medium composition is described in detail in (20). The pH was monitored using a pH-sensor EasyFerm Plus (Hamilton, Reno, NV, USA). During cultivations, pH was kept constant at 5.0 and was controlled with base addition only (12.5% NH_4_OH), while acid (10% H_3_PO_4_) was added manually, if necessary. The temperature was kept constant at 30 °C. Aeration was carried out using a mixture of pressurized air and pure oxygen at two vvm to keep dissolved oxygen (dO_2_) above 30% at all times. The dissolved oxygen was monitored using a fluorescence dissolved oxygen electrode, Visiferm DO (Hamilton, Reno, NV, USA). After sugar during batch phase was depleted, the continuous operation was started. During chemostat cultivation volume in the reactor was adjusted and maintained constant via an immersion tube connected to a bleed pump.

Samples were taken during chemostat cultivations at alternating time-points depending on the morphological alterations. These alterations highly depend on the cultivation time and the set dilution rate, with details given here (21). Microscopic analysis was used to observe pseudohyphae growth.

##### A.1.4 Cultivation of *K. phaffii CBS7435*

*K. phaffii CBS7435, △DAS1, △DAS2, △AOX1* was cultivated in M2 citrate-buffered media (22) supplemented with 20 mg/L thiamine pyrophosphate (TPP). A single yeast colony from a YPD (1% yeast extract, 2% peptone, 2% glucose, and 1.5% bacteriological agar) agar plate was inoculated into a 250 mL shake flask containing 50 mL of YPD broth as a preculture grown at 28 °C and 200 rpm. The samples for microscopy were taken from cultures grown in M2 citrate-buffered media with 1% (v/v) methanol as the carbon source. The dataset from this cultivation will be referred to as *Komagataella phaffii* strain set_A in this article.

##### A.1.5 Generation of *A. niger* conidia

An industrial strain of *A. niger* was spotted on minimal medium plates from a -80 °C glycerol stock. The plates were incubated at 30 °C for 7 days to allow for sufficient conidiation. Conidia were harvested by pouring 10 mL of 0.1% Tween-20 solution onto the plates and were subsequently scraped off by using a sterile cotton swab. The suspension was filtered using sterile Miracloth to remove agar debris and mycelium. After filtering the suspension was centrifuged at 3000g for 15 minutes at 4 °C. Supernatant was discarded and the centrifugation step was repeated after 10 mL of 0.1% Tween-20 solution was added. After centrifugation, the supernatant was discarded, and up to 10 mL of 0.1% Tween-20 solution was added.

#### A.2 Microscopy and Imaging

Unstained wet mounts of the *A. pullulans* strains, *K. phaffii* (strain set_A), *A. niger* spores, and *T. reesei* were prepared and observed using brightfield microscopy. For all *A. pullulans* strains, microscopic imaging was performed at multiple time points ranging from 24h to 240h. Images were captured using a Nikon ECLIPSE E200 light microscope, equipped with a BRESSER Mikrookular CMOS sensor-based Full HD eyepiece camera (Art. No. 5913650). Images of *T. reesei, K. phaffi* (strain set_A) and *A. niger* were captured using the same setup but with magnification of 100x and 200x for *T. reesei,* and 100x, 200x and 400x for *K. phaffi* (strain set_A). Appropriate dilutions of the *A. niger* spore suspension were prepared in 0.1% Tween-20 solution, and the conidia were observed using an improved Neubauer counting chamber at 200x magnification. Image acquisition and export was performed using the “Breeser CamLab Lite” software at a resolution of 1920 x 1080 (width x height).

Microscopic analysis of *K. phaffii* GS115 was performed using an Olympus CKX41 inverted microscope (Olympus Life Science, Tokyo, Japan) with an IX2-SLP phase contrast slider (Olympus Life Science, Tokyo, Japan) using a Canon EOS 250D (Canon, Tokyo, Japan) camera.

Microscopy and imaging of *A. melanogenum* was performed using a Leica DMi8 microscope. Images were captured at multiple time points ranging from 24h to 168h, at 100x, 400x, and 630x magnification. Images with a resolution of 2560 x 1920 (width x height) were exported using Leica Application Suite.

##### A.2.1 Dataset expansion strategy

We applied a dataset expansion strategy in which the 1920×1080 pixel images were cropped into 640×640 pixel crops, with each representing a unique image (Figure 1). This was inspired by a similar strategy applied by Li et al. (11) & Juneja et al. (23), which led to an increase in the number of images available for the dataset and more efficient training. A custom Python script “cropper.py” was used to generate six unique images of size 640×640 from a single 1920×1080 image. The script was also designed to handle the edge cases, where the region of the crop is adjusted if the crop section is out of bounds of the original image size. 15437 images were generated using this dataset expansion strategy.

**Figure 1:**
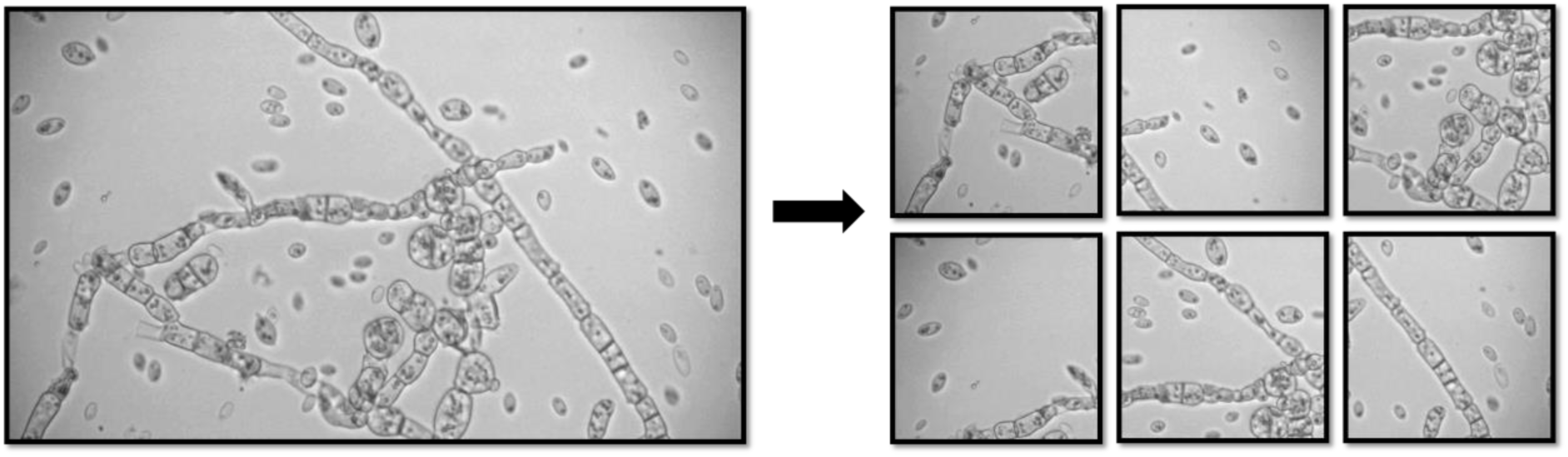
Representative full-size (1920 x 1080 pixels) microscopic image of A.pullulans (left panel) cropped into six unique 640 x 640 pixel crops (right panel) as part of the dataset expansion strategy.

#### A.2.2 Image annotation

Images with 640X640 resolution were then manually screened and annotated using LabelMe (v5.4.1) (24). Objects were annotated with rectangular bounding boxes. In order to track the number of objects from each class, for monitoring and class balancing operations. A Python script, “annotation handler.py,” was used in our study.

The Python library Labelme2YOLO (v0.1.7) and Labelme2YOLO (v0.2.5) were used to convert annotations in “.JSON” format to “YOLO” annotation format (25). The dataset was auto-split into an 80:20 Train: Validation split during the conversion.

#### A.2.3 Training and Validation datasets

The dataset generation was an iterative process involving repeated manual review of the datasets, removal of noisy or incorrect labels, and addition of new images to the datasets. This led to the creation of three versions of the dataset: X-ray, Yankee, and Zulu. Zulu was the final version of the dataset. X-ray, Yankee, and Zulu datasets consisted of 1028 images, 1002 images, and 1504 images with 8575, 8652, and 11804 objects, respectively, belonging to 13 different classes. See “S2 Table 2” for detailed information on the distribution of objects across all the classes in training and validation sets, in all five splits, and three dataset families.

#### A.2.4 Test dataset

We created an independent global test dataset of 166 images from our primary image bank, which are not part of any of the X-ray, Yankee, and Zulu datasets. The test dataset contained images of *A. pullulans, A. melanogenum*, and *T. reesei*. See “S3 Table 3” for class-wise object distribution in global test dataset. This test dataset was used to test all model families using the validator function built into the Ultralytics framework (26).

#### A.2.5 5-fold cross-validation

To make the optimum use of our datasets, we performed K-fold stratified cross-validation using a Python cross-validation split script by (27) as the base script and modified it. With the script “kfold_splitter.py,” every dataset family was split into five folds in order to achieve a balance between class representation and computational costs. A stratified k-fold cross-validation was preferred over a simple k-fold cross-validation in order to account for the imbalanced nature of the dataset (28). All the computations were performed on Vienna Scientific Cluster-5, powered with an NVIDIA A100 GPU.

#### A.2.6 Auto-labelling

The final version of the dataset, Zulu, was a hybrid dataset containing auto-labelled images generated with the Yankee_s5 version of our model trained on pre-trained YOLOv11m. The auto-labeling was performed on images from our dataset, which were not part of any previous dataset. A Python script, “datafeed_prediction.py,” was used to run the prediction function of the model on the images. The prediction function outputs the images with annotated objects in bounding boxes, along with corresponding text files with annotations in YOLO format. Post prediction, all 14200 images were manually reviewed to check for correct annotations. Five hundred seventeen images were selected and then added to the Yankee dataset images. The dataset was manually reviewed in order to remove any identical images. This resulted in the final dataset with 1504 images.

### B. Training TU_MyCo-Vision on Ultralytics YOLO with custom dataset using transfer learning approach

#### B.1.1 Initial training and hyperparameter exploration

Initially, the pre-trained YOLOv8x and, after its release, YOLOv11x and YOLOv11m models were used and trained on our custom dataset using transfer learning. We randomly selected x-ray_split_1 as the representative dataset among the five dataset splits. Keeping the default Ultralytics training hyperparameters as a reference, we arrived at the list of hyperparameters as shown in “S4 Table 4” through exploratory trials. A detailed list of all the default hyperparameters and augmentation settings can be found at (29).

Throughout the study, we considered a mAP@50-95 > 0.5 for single-cell classes and mAP@50 > 0.5 for filamentous classes as the criteria for a well-performing model. Moreover, validation metrics like class loss, box loss, and distribution focal loss (dfl) were monitored to prevent overfitting or any instabilities in training. All metrics and loss functions were generated by the Ultralytics framework.

#### B.1.2 Hyperparameter optimization

Based on the hyperparameters obtained after the exploratory search, a hyperparameter search was performed. These parameters (refer to “S4 Table 4”) were used to define the search space for the hyperparameter tuning. For tuning the hyperparameters of the YOLOv8x model on our custom dataset, the “model.tune()” method was used. This utilized the “Tuner” class based on the Ultralytics YOLO mutation algorithm to search for the best-fit hyperparameters (30). Atotal of 45 iterations were executed, each with a limit of 300 epochs, using the Stochastic Gradient Descent (SGD) optimizer. During each iteration, the current parameter set was mutated, and models were evaluated using the fitness score, calculated based on the key performance indicators.

#### B.2 Cross-validation runs

Using the tuned hyperparameters (see “S5 Table 5”), we performed the training with all five splits obtained after the 5-fold stratified dataset splitting. This process was performed for all the models except for the model trained on the dataset Zulu. For training the Zulu model family, augmentation parameters were slightly modified for more robust augmentation. Refer to “S6 Table 6” for the list of hyperparameters used for model Zulu.

#### B.3 Model Evaluation

The object detection model was evaluated using Intersection over Union (IoU), box-level precision (P) and recall (R), and mean average precision (mAP). The mAP was computed at an IoU threshold of 0.50 (mAP@50) and also averaged across thresholds from 0.50 to 0.95 in 0.05 increments (mAP@50–95) to assess the performance across varying degrees of overlap.

During model training, three loss functions were used to guide the learning process: classification loss, box loss, and distribution focal loss (dfl). Classification loss was evaluated to assess the performance of the model in correctly classifying objects of different classes. A lower classification loss indicates better performance in distinguishing between different cell morphologies. Bounding box loss penalizes spatial misalignment between predicted and ground truth boxes. Distribution focal loss (dfl) was used to improve the precision of bounding box predictions by focusing learning on difficult-to-detect objects.

The Ultralytics framework also generated visual outputs to support the interpretation of the model’s performance. These included F1 score curves, precision-recall (PR) curves, along with raw and normalized confusion matrices. Qualitative assessments involved visualization of annotated validation images, comparing ground truth and predicted bounding boxes. All results, including numerical metrics and visual outputs, were saved in the runs/detect/val directory for further analysis (31)

### C. Graphical User Interface Implementation and Data Analysis Workflow

To improve user accessibility by eliminating the need to install dependencies or run the Ultralytics framework via terminal, a cross-platform graphical user interface (GUI) was developed in Python (version 3.10 or higher) using multiple open-source libraries. The interface was implemented using customtkinter (v5.2.2) (32), a wrapper around the standard tkinter library (33). All image handling operations were performed using Pillow (v11.2.1) (34), and OpenCV (35), including image dimension capture, resizing, and rendering. The prediction script was integrated into the GUI, allowing parameterization through interface controls. Except for the imgsz parameter (automatically determined from image dimensions) and interface-based inputs, all model parameters were kept at default values.

Annotation data was extracted and analyzed for quantitative assessment of predicted morphotypes.

Edge Bounding Box Exclusion: A key data preprocessing step involved the exclusion of bounding boxes positioned at the edges of images to avoid counting partially visible cells (false positives or incorrect detections). This feature was disabled for hyphal morphologies and clumps (C7-E, C7-F, C7-G, and C6), as they typically occupy large regions in the image, resulting in bounding boxes that extend to the image edges. Bounding boxes were excluded from the analysis if either the left or right edges (calculated as center x ± width/2), or the top or bottom edges (center y ± height/2), were within a small threshold (epsilon = 0.001) of the image border (0 or 1 in normalized coordinates).

Class Exclusion: To enable targeted analysis, a class exclusion logic was implemented whereby selected morphotypes could be grouped into an “Others” category during processing. Activating this feature resulted in the export of two data tables: a filtered data table (filename_excluded.csv) and a raw data table (filename_rawdata.csv). This was tested using a subset of the test dataset by grouping filamentous morphologies (C7-E, C7-F, C7-G) into the “Others” category for single-group analysis.

Single and Multi-Group Analysis: For single-group analysis, a random subset of 90 images from the global test set was analyzed using the single-group pipeline, with and without class exclusion enabled. For multi-group analysis, a test set was constructed with 10 *C. albicans* images (Candescence dataset) (8) 9 *K. phaffii (strain set_A)*, 9 *K. phaffii GS115*, and 9 *A. niger* spore images. Predictions were performed using default settings, and results were organized into separate folders representing each group. The multi-group analysis pipeline was then applied to compare morphological profiles across these groups.

The data analysis pipeline generated interactive visualizations, including absolute count bar plots, relative abundance bar plots, mean relative abundance bar plots, normalized stacked bar plots, and hierarchical clustering heatmaps using Ward’s method. Hierarchical clustering was performed using the scipy.cluster.hierarchy module in the SciPy library (36). For multi-group analysis, an additional heatmap comparing all individual images across groups was also generated. All plots were rendered interactively via the Plotly library (v6.1.2) (37) using the generated “.csv” files as input.

## Results

### A. Characterizing morphological variation in *A. pullulans*

After the cultivation of twelve *A. pullulans* strains in ten different cultivation media, we classified the cell morphologies into thirteen different classes, based on phenotypic observations. These morphologies were broadly categorized into single-celled morphologies, filamentous morphologies, and special morphologies. C1, C2, C2-B, C3-B, C5, and C9 are single-cell classes. C7-G, C7-E, and C7-F are classes for filamentous morphologies, and C4, C6, and C11 are special classes representing budding events, clumping, and irregularly shaped cells, respectively (See Figure 2). Refer to Table 1 for a detailed description of object classes.

**Figure 2:**
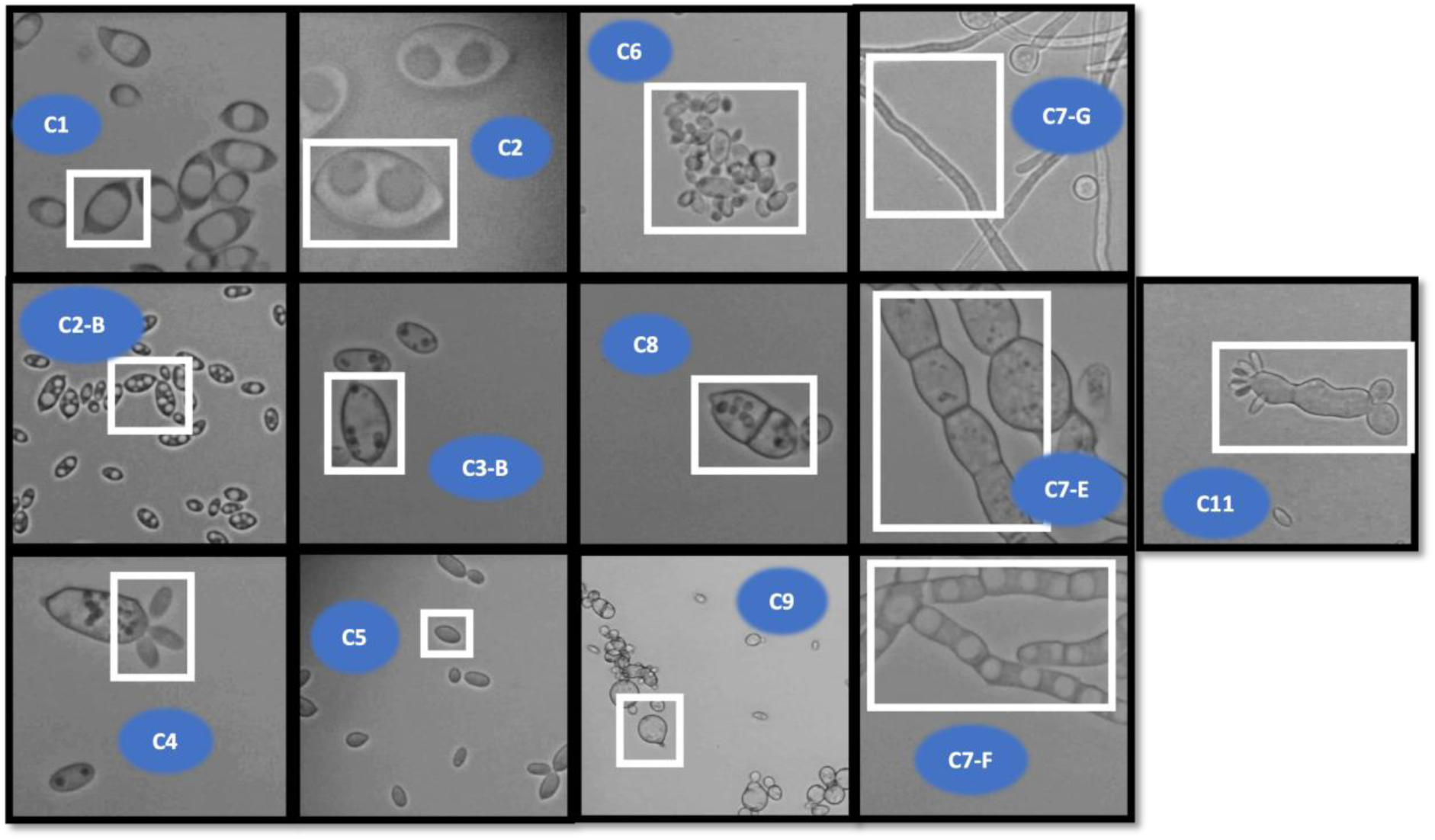
TU_MyCo-Vision classes and representative cell morphologies, highlighted with white rectangular boxes. C1, C2, and C2-B are vacuolated cells. C3-B are cells with granular cytoplasm. C5 are general yeast-like cells lacking any special features. C9 cells are spheroidal-shaped cells. C4, C6, and C11 are special cell types representing budding events, cell clumping/aggregates, and irregular cell shapes. C7-E (septate hyphae/ pseudo hyphae), C7-G (smooth continuous filamentous/true hyphae), and C7-F (vacuolated hyphae) are filamentous cell classes.

**Table 1:** Detailed description of object classes in TU_MyCo-Vision

| Class | Cell Shape | Description |
| --- | --- | --- |
| C1 | Ovoid, oblong, spheroidal, elongated | Eye-like or egg-like appearance due to the presence of a single vacuole-like structure |
| C2 | Ovoid, oblong, spheroidal, elongated | Binocular appearance due to dual vacuole-like structures |
| C2-B | Ovoid, oblong, spheroidal, elongated | Multiple vacuole-like structures |
| C3-B | Ovoid, oblong, spheroidal, elongated | The granular appearance of the cytoplasm due to small bead-like structures |
| C4 | NA | Indicates budding events |
| C5 | Ovoid, oblong, elongated | Lacks any distinctive morphological features. Includes all yeast like cells |
| C6 | NA | Cell aggregates and clumping |
| C8 | NA | Septate cells |
| C9 | Spheroidal | Spherical/Spheroidal cells without distinctive morphological features |
| C11 | NA | Irregular-shaped cells that cannot be categorized in the other classes |
| C7-E | Filamentous | Filaments with an appearance similar to septate hyphae |
| C7-F | Filamentous | Filaments with vacuole-like structures |
| C7-G | Filamentous | Smooth filaments without any distinctive features |

### B. Initial training, Hyperparameter exploration, and optimization

Initial training was conducted with the YOLOv8x architecture using the split_1 dataset of X-ray family, selected as a representative subset from the stratified 5-fold cross-validation splits. Using default Ultralytics hyperparameters as a baseline, an exploratory approach was employed to identify a reasonable set of hyperparameters (see “S4 Table 4”), which yielded satisfactory early-stage performance across several morphological classes.

Version 1 of the model demonstrated high detection accuracy for major single-cell classes. Specifically, mAP@50 values were 0.840 for class C1 (cells with central vacuole), 0.816 for C2 (cells with dual vacuoles), 0.785 for C5 (yeast-like morphologies), 0.822 for C2-B (multiple vacuoles), 0.904 for C3-B (granular cytoplasm), and 0.890 for C8 (septate cells). In contrast, filamentous morphologies posed greater challenges: the model achieved mAP@50 values of 0.811, 0.566, and 0.547 for C7-E (septate hyphae or pseudohyphae), C7-F (vacuolated hyphae), and C7-G (smooth hyphae/nonseptate hyphae), respectively.

Evaluation of the normalized confusion matrix revealed misclassification events. Class C5 (yeast-like cells) was particularly prone to confusion, with frequent misclassification as C11 (irregular shapes) and C9 (spheroidal cells). Although C5 achieved 80% correct predictions, it was also associated with a hallucination rate (i.e., proportion of predicted objects of a class that do not correspond to any ground truth instances of that class) of 36%. These findings highlight the morphological overlap among spheroidal and ovoid classes, highlighting a key challenge in differentiating single-cell morphologies with geometrical overlap.

The highest mAP@50-95, of 0.5216, was achieved at epoch 486. This level of performance met our minimum criteria for a functional prototype, and the hyperparameter set was selected as a baseline for hyperparameter optimization. Refer to “S7 Folder (V1)” for PR curves, F1 plot, confusion matrices, and other result visualizations.

Subsequent optimization was conducted using Ultralytics genetic mutation-based tuner, which adapts hyperparameters iteratively to maximize fitness (30).

The best-performing configuration emerged at iteration 19, with a fitness score of 0.5491. This iteration produced a model with a precision of 73.21%, a recall of 68.01%, a mAP@50 of 0.7268, and mAP@50–95 of 0.5293. These results marked an improvement over the baseline exploratory parameters. Notably, fitness scores plateaued after iteration 19, with the second-best score (0.5404) obtained at iteration 23 (Figure 3), indicating diminishing returns from further mutation. The hyperparameters derived from the optimal iteration (listed in “S5 Table 5”) were selected for use in all subsequent training across the dataset splits. This tuning process contributed to performance consistency and stability in downstream model families (X-ray, Yankee, and Zulu), particularly in reducing false positives and improving the balance between sensitivity and specificity. These results validate the effectiveness of targeted hyperparameter optimization in enhancing detection for diverse fungal cell morphologies, especially when addressing under-represented and structurally complex classes such as filamentous forms. Refer to “S8 Folder “hyperparameter_tuning” for additional information.

**Figure 3.**
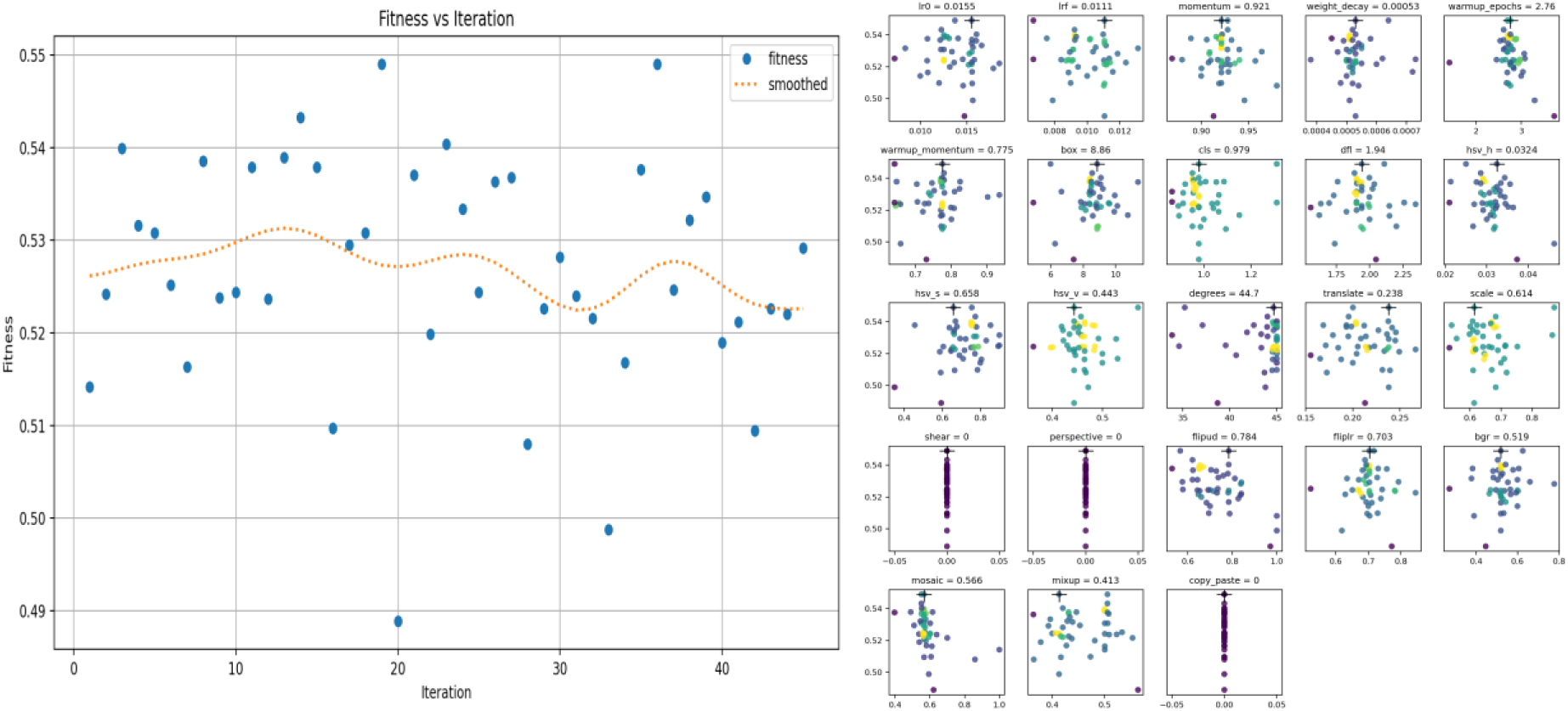
(Left Panel) Fitness vs. iteration plot. Fitness scores across 45 iterations of hyperparameter tuning. The dotted line indicates a smoothed trend. Scatter plots of tuned hyperparameters vs. fitness. (Right Panel) Each subplot shows the relationship between the hyperparameter and resulting fitness scores during the hyperparameter evolution. Annotated values indicate the best-performing parameters. Parameters set to ‘0’ were not tuned during the hyperparameter evolution.

### C. k-fold cross-validation runs

Next, k-fold cross-validation training was performed, initially with the X-ray and Yankee dataset, and finally with the Zulu dataset. Even though the initial choice of model was YOLOv8x. We migrated to the newer model YOLOv11x and later YOLOv11m. The YOLOv11x to YOLOv11m was performed because YOLOv11m is a smaller model, which is computationally more efficient and reduces overfitting, especially on smaller datasets (38). The evaluation of all the models was performed using the standard object detection metrics, including box precision, box recall, mAP@50 and mAP@50-95.

#### C.1 Performance evaluation of X-ray, Yankee, and Zulu models on validation dataset

We trained and evaluated three model families, X-ray, Yankee, and Zulu, each corresponding to a new improved version of the TU_MyCo-Vision dataset. All models were trained using the five dataset splits generated after 5-fold cross-validation and assessed based on standard object detection metrics (See Table 2 for performance summary of three model families). In each family, the dataset split that resulted in the best model performance was considered as the representative model and was further analyzed using confusion matrices, precision-recall (PR) curves, and training/validation loss curves.

**Table 2:** Comparative summary of average performance metrics across five dataset splits of X-ray, Yankee, and Zulu model families evaluated on the validation dataset

| Metric | X-ray | Yankee | Zulu |
| --- | --- | --- | --- |
| Dataset version (No. of images) | 1028 | 1002 | 1504 (Manual + auto labelled) |
| Architecture | YOLOv8x | YOLOv11m | YOLOv11m |
| Best Split | xray_split4 | yankee_split5 | zulu_split1 |
| Highest mAP@50-95 at epoch | 309 | 405 | 483 |
| Precision (mean ± SD) | 71.56 ± 0.0174 | 72.43 ± 0.0160 | 74.59 ± 0.0230 |
| Recall (mean ± SD) | 71.83 ± 0.0356 | 71.67 ± 0.0203 | 72.97 ± 0.0163 |
| mAP@50 (mean ± SD) | 0.7443 ± 0.0202 | 0.7535 ± 0.0147 | 0.7745 ± 0.0106 |
| mAP@50-95 (mean ± SD) | 0.5360 ± 0.0127 | 0.5357 ± 0.0096 | 0.5732 ± 0.0123 |

The X-ray split_4 model achieved high classification accuracy (i.e., all detected objects were assigned to their correct morphological class, according to the normalized confusion matrix) for common morphotypes, including C1 (89%), C2 (88%), C5 (84%), C6 (90%), C8 (88%), and C2-B (79%), with moderate performance for C3-B (76%), C4 (73%), and C11 (58%). C9 was frequently misclassified as C5 (21%), and misclassification of C1 primarily occurred with C2 (4%) and C2-B (6%). Filamentous classes were particularly challenging: C7-E, C7-F, and C7-G were correctly predicted in 75%, 58%, and 52% cases, respectively, with 20-45% of bounding boxes assigned to background or neighboring filamentous classes. Based on the PR curve, corresponding mAP@50 values were 0.649, 0.440, and 0.429, respectively, while higher scores were observed for C1 (0.914), C2 (0.902), C2-B (0.939), C6 (0.921), and C8 (0.882).

The Yankee split_5 model achieved greater than 80% accuracy for C1, C2, C3-B, C5, C6, C8, and C7-E. Detection rates for C7-F and C7-G were 55% and 51%, respectively. Misclassifications were prominent between C9 and C5 (18%), C3-B and C5 (14%), and C8 and C11 (10%). PR curve analysis showed mAP@50 values of 0.934 (C1), 0.899 (C2-B), 0.914 (C8), 0.854 (C3-B), 0.756 (C7-E), 0.592 (C7-F), and 0.500 (C7-G). Lower mAP@50 scores were also noted for C4 (0.633), C9 (0.699), and C11 (0.634).

The Zulu split_1 model demonstrated high classification accuracy across several classes: 95% for C1, 89% for C6, 84% for C3-B, and less than or equal to 80% for C2, C5, and C2-B. C4 achieved 69% classification accuracy, although 25% of its instances were missed. C11 remained difficult to classify, with 50% classification accuracy and frequent misclassification into C6, C9, or background. The detection of filamentous morphotypes improved, with C7-E, C7-F, and C7-G achieving 76%, 71%, and 67% classification accuracy, respectively, and corresponding mAP@50 values of 0.808, 0.555, and 0.625. C4 and C11 recorded mAP@50 scores of 0.753 and 0.681, respectively, based on the PR curve analysis.

Overall, all three model families exhibited stable training dynamics, with steadily decreasing loss curves and consistent improvements in all metrics over epochs. Successive models benefited from dataset enhancements, which contributed to improved performance metrics, particularly for morphologically complex or rare cell types. Among the three, Zulu demonstrated superior performance to its predecessors and showed reliable performance for both single-celled morphologies and complex filamentous classes, supporting its deployment as the final representative model for TU_MyCo-Vision. Refer to the supporting information folder “S9 xray”,” S10 zulu”, and “S11 yankee” for detailed metrics, confusion matrix, PR curve, F1 curve, and other visualizations.

### D. Comparison of all TU_MycoVision model families on the global test dataset

A separate global test dataset of 166 images was created in order to test all the models. The test set was curated manually and contained images that are not part of any of the datasets used for the training of our 3 model families. Using the global test dataset, we compared all the models belonging to the X-ray, Yankee, and Zulu model families.

15 models were evaluated based primarily on mAP@50-95, precision, and recall performance, with more preference to mAP@50-95 and Precision metrics in the same order of priority for selecting the best performing model. Out of which Zulu_s3 was the best performing model, with a high precision of 73.4%, a recall of 66.5%, and mAP@50-95 of 0.545. These improvements could be attributed to the consistent improvement of our datasets over each family of models, X-ray, Yankee, and Zulu. Table 3 describes the detailed performance metrics of all the models. Zulu_s5 and zulu_s4 also showed robust performance with mAP@50-95 of 0.546 and 0.527, respectively. Relatively lower mAP@50-95 scores were observed for X-ray models, ranging from 0.493 to 0.513, and also for Yankee models, whose scores ranged from 0.498 to 0.526. This makes the Zulu family of models the best models with a mAP@50-95 ranging from 0.519 to 0.546 to detect 13 fungal cell morphologies in microscopic images.

**Table 3:**
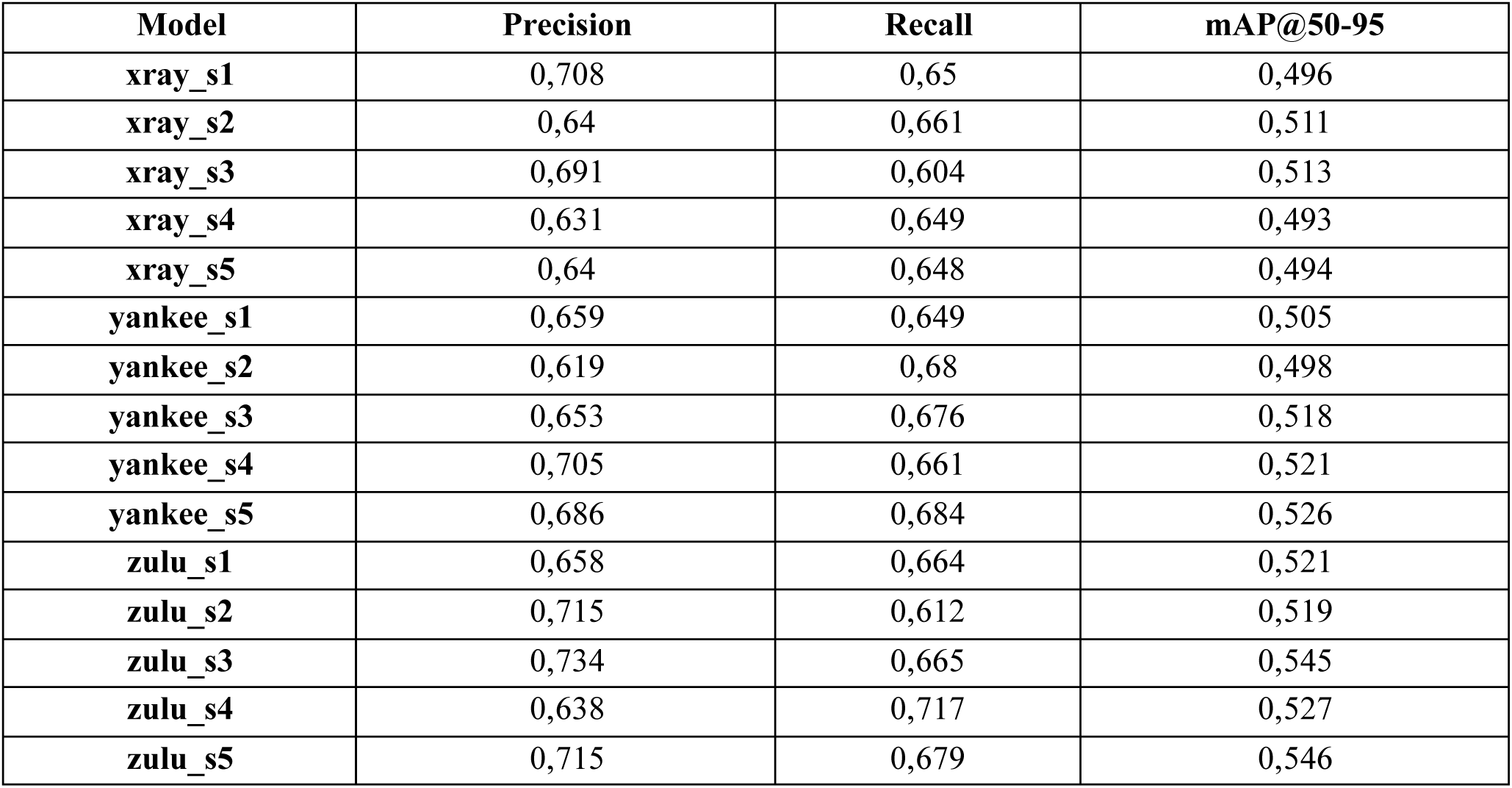
Precision, Recall, and mAP@50-95 performance comparison of all TU_MyCo-Vision model families on the global test dataset

| Model | Precision | Recall | mAP@50-95 |
| --- | --- | --- | --- |
| xray_s1 | 0,708 | 0,65 | 0,496 |
| xray_s2 | 0,64 | 0,661 | 0,511 |
| xray_s3 | 0,691 | 0,604 | 0,513 |
| xray_s4 | 0,631 | 0,649 | 0,493 |
| xray_s5 | 0,64 | 0,648 | 0,494 |
| yankee_s1 | 0,659 | 0,649 | 0,505 |
| yankee_s2 | 0,619 | 0,68 | 0,498 |
| yankee_s3 | 0,653 | 0,676 | 0,518 |
| yankee_s4 | 0,705 | 0,661 | 0,521 |
| yankee_s5 | 0,686 | 0,684 | 0,526 |
| zulu_s1 | 0,658 | 0,664 | 0,521 |
| zulu_s2 | 0,715 | 0,612 | 0,519 |
| zulu_s3 | 0,734 | 0,665 | 0,545 |
| zulu_s4 | 0,638 | 0,717 | 0,527 |
| zulu_s5 | 0,715 | 0,679 | 0,546 |

Considering zulu_s3 as our best model, we further analyzed the PR curves and normalized confusion matrix to get more performance insights. The precision recall curve indicates strong performance for single cell classes like C2, C5, C1, C2-B, C3-B, and C8 with an average precision at (50% IoU) of 91.2%, 79.25%, 84.4%, 88.7%, 90%, and 89.7% respectively, based on PR curve analysis (Figure 4). The model performed best for C1, C2, C2-B cell types, which represent vacuolated morphologies, and C3-B, which represents single cells with granules in the cytoplasm, and C8, which are septate cells. Satisfactory performance was observed for the classes C4 (budding events), C6 (cell aggregates/clumps), and C11 (irregular morphologies).

**Figure 4.**
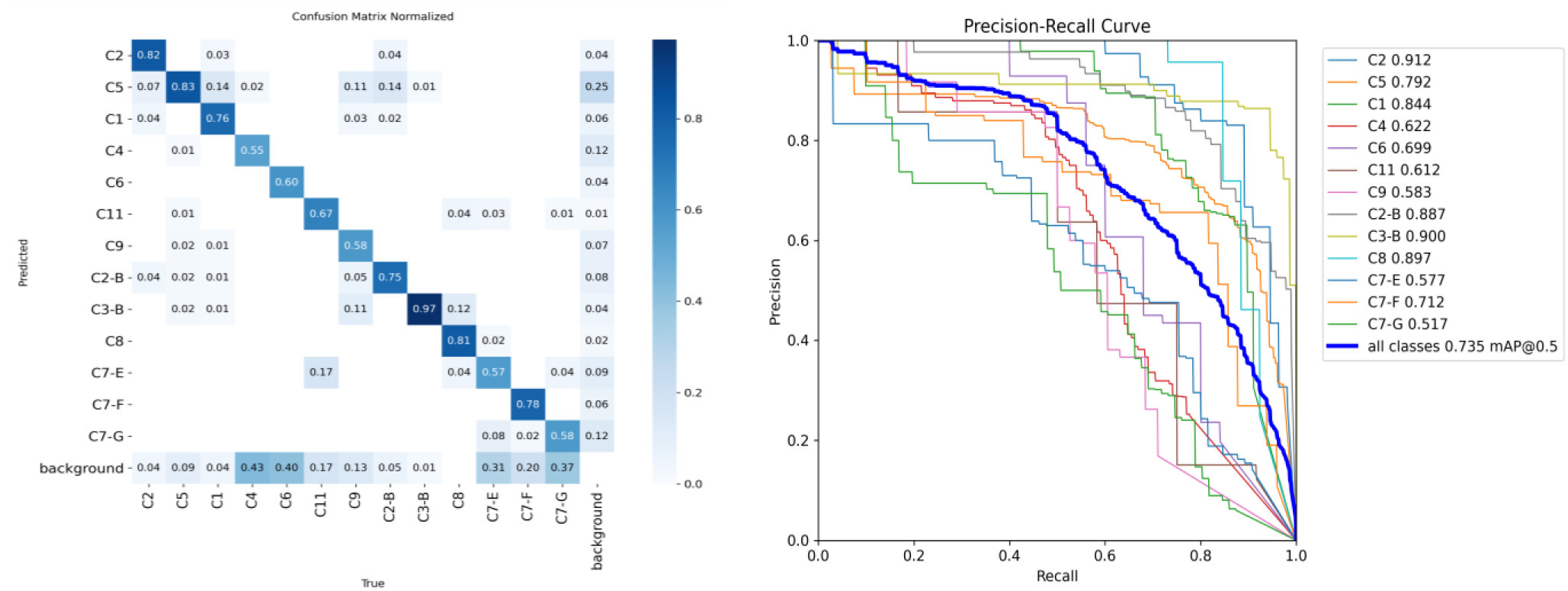
Normalized confusion matrix (left) of the Zulu_s3 model validated on the global test set. Precision-Recall Curve (right) of the zulu_s3 model validated on the global test set.

The model also performed well for the filamentous cell classes with average classification accuracy values consistently above 51% based on the normalized confusion matrix (Figure 4), which is satisfactory for such complex morphologies. Overall, our model demonstrated detection accuracy of 73.5% for 13 different cell morphologies (Refer Table 4 for detailed class-wise performance metrics of model zulu_s3). The normalized confusion matrix provides deeper insights into the model’s misclassifications (Figure 4). High classification performance was observed for the classes C2 (82% correct predictions), C5 (83% correct predictions), C3 -B (97% correct predictions), and C8 (81% correct predictions). Misclassification was particularly observed between classes C1:C5 with 14% C1 cells misclassified as C5, C5:C9 (11% C9 misclassified as C5), C9:C3-B (11% C9 misclassified as C3-B), C2-B:C5 (14% C2-B misclassified as C5), C8:C3-B (12% C8 misclassified as C3-B). Satisfactory performance was observed for the filamentous classes, with classification accuracies of 57% for C7-E (septate hyphae/pseudohyphae), 78% for C7-F (vacuolated hyphae), and 58% for C7-G (smooth hyphae). Although the numbers for the filamentous classes seem to be on the lower side, it has to be considered that the hyphal structures are mostly spread across large areas along with their complex structures, which affects their detection.

**Table 4:** Class-wise performance metrics of the zulu_s3 model evaluated on the global test dataset

| zulu_s3 | Class | Images | Instances | Precision | Recall | mAP@50 | mAP@50-95 |
| --- | --- | --- | --- | --- | --- | --- | --- |
|  | **all** | 166 | 956 | 0.734 | 0.665 | 0.735 | 0.545 |
|  | **C2** | 27 | 55 | 0.839 | 0.851 | 0.912 | 0.730 |
|  | **C5** | 77 | 229 | 0.709 | 0.789 | 0.792 | 0.587 |
|  | **C1** | 31 | 78 | 0.762 | 0.731 | 0.844 | 0.677 |
|  | **C4** | 63 | 139 | 0.811 | 0.475 | 0.622 | 0.327 |
|  | **C6** | 16 | 25 | 0.639 | 0.600 | 0.699 | 0.560 |
|  | **C11** | 8 | 12 | 0.613 | 0.583 | 0.612 | 0.480 |
|  | **C9** | 27 | 38 | 0.557 | 0.579 | 0.583 | 0.459 |
|  | **C2-B** | 28 | 95 | 0.865 | 0.739 | 0.887 | 0.729 |
|  | **C3-B** | 36 | 74 | 0.778 | 0.949 | 0.900 | 0.730 |
|  | **C8** | 20 | 26 | 0.900 | 0.846 | 0.897 | 0.790 |
|  | **C7-E** | 36 | 65 | 0.664 | 0.446 | 0.577 | 0.338 |
|  | **C7-F** | 32 | 49 | 0.720 | 0.571 | 0.712 | 0.391 |
|  | **C7-G** | 30 | 71 | 0.680 | 0.479 | 0.517 | 0.291 |

See the supporting folder “S13 validation_global_set” for validation performance visualizations of all models validated on the global test dataset.

Refer to the supporting information table “S14 Table 7. validation_metrics_global_test_set” for detailed validation metrics of all models.

### E. Graphical User Interface, and Data Analysis

We developed a cross-platform graphical user interface to make the detection tool accessible and user-friendly. As all the dependencies and libraries are packaged into a single executable “.exe” for Windows-based systems and “.app” for macOS, the need for the user to set up the required environment is eliminated. This makes the tool suitable for users without advanced computational skills. The GUI is organized into three main panels: Detection, Analysis, and Results Dashboard. Detection panel includes all options from selection of I/O folders, class selection, model parameter adjustments, and prediction execution (Figure 5a). The Analysis panel allows the user to perform single-group and multi-group analyses, along with the class exclusion panel for selecting morphological classes to be grouped as “others” (Figure 5c). The Results Dashboard offers an interactive side-by-side visualization of original and processed images for easy visualization of results (Figure 5b). Quantitative results are accessible via an embedded CSV table, which can be loaded and viewed in the Results Dashboard (Figure 5d). The tool, using the loaded CSV file, also generates and renders plots after clicking the “Click to View Plots” button.

**Figure 5a.**
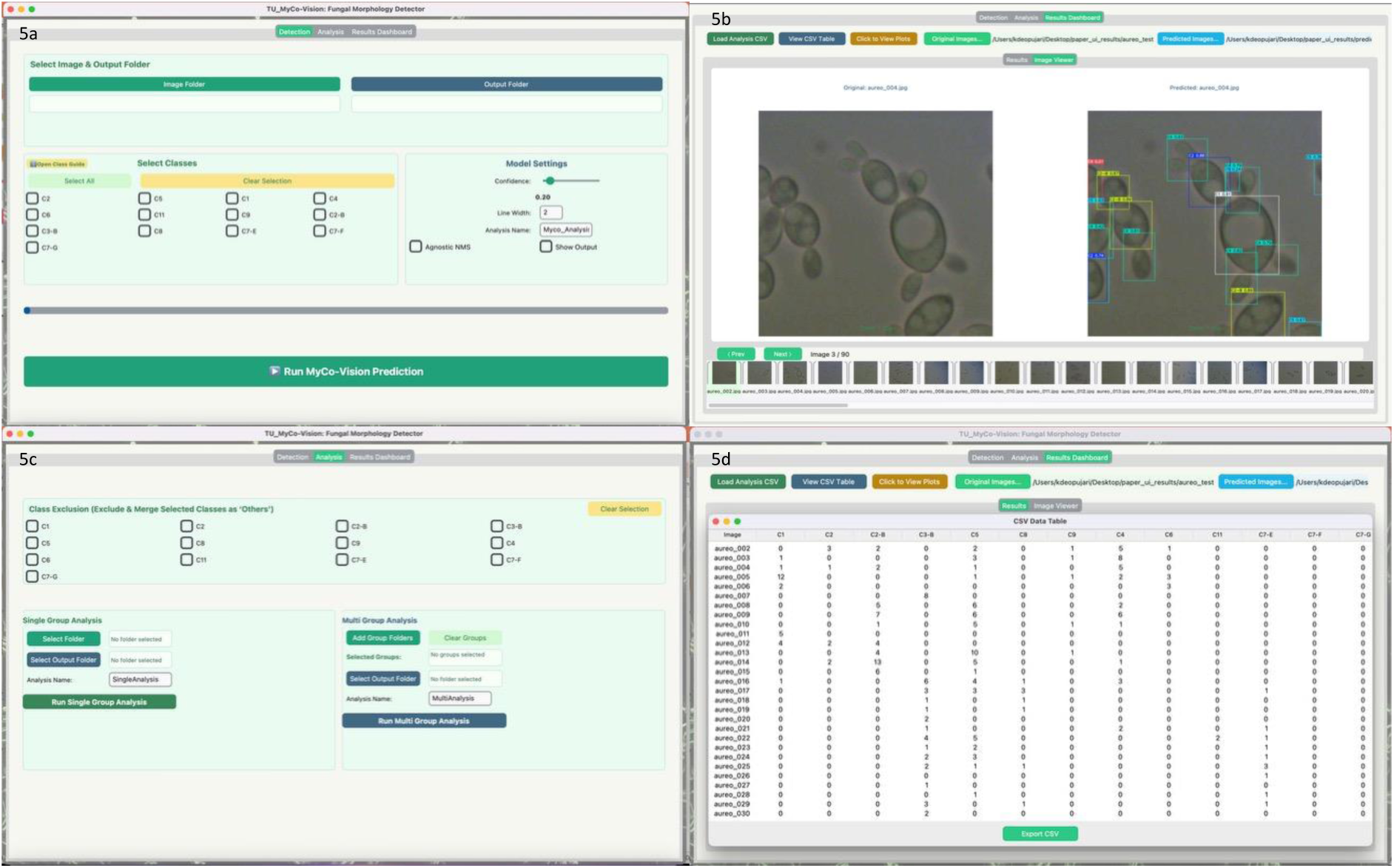
Detection panel, enabling users to select input/output folders, select morphological classes, adjust model parameters, and execute prediction. Figure 5b. Result Dashboard panel image viewer displaying paired original and annotated images side-by-side, with navigation panel. Figure 5c. Analysis panel, which includes the class exclusion feature and options for single-group and multi-group analysis, along with the option to select input/output folders and analysis name. Figure 5d. Results Dashboard panel CSV data table view.

#### E.1 Single-Group Analysis of Test Set Validates the Data Processing and Visualization Pipeline

We tested the data analysis and visualization pipeline for the single-group analysis function using 90 images from our global test set. Both workflows were evaluated: one with all classes and another with class exclusion activated, in which all filamentous classes (C7-E, C7-F, and C7-G) were grouped together into the “Others” category. The pipeline processed all 90 test images and produced six quantitative visualizations, including absolute count bar plots, normalized stacked bar plots, relative and mean relative abundance plots, and heatmaps with hierarchical clustering (Figures 6 and 7). This demonstrated the tool’s ability to handle a large number of images, apply class-exclusion logic, and generate quantitative visualization plots within a single, integrated environment.

**Figure 6a:**
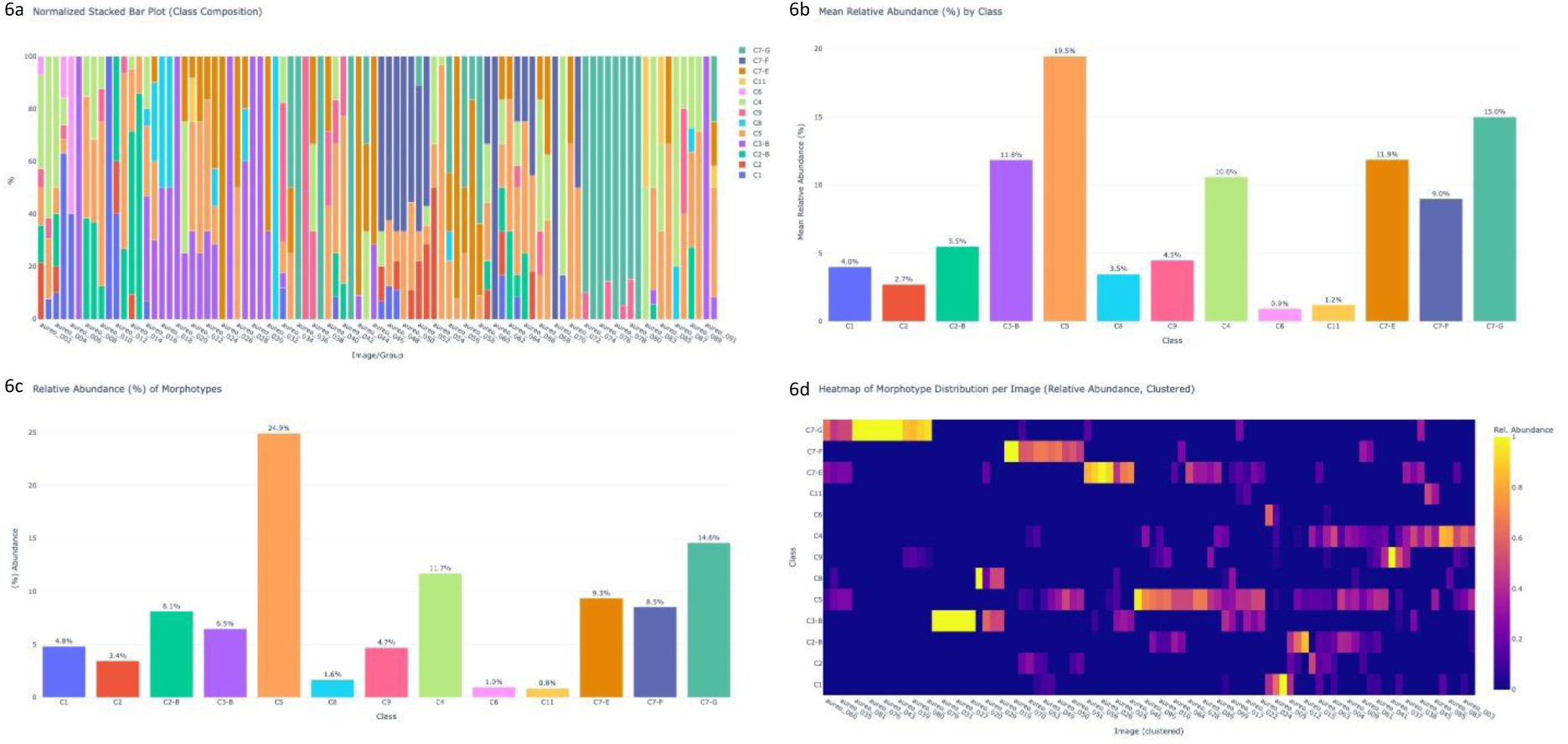
Normalized stacked bar plot representing morphological composition across 90 test set images. Figure 6b: Mean relative abundance (%) of each morphological class across all images. Figure 6c: Relative abundance (%) of each morphological class across the dataset. Figure 6d: Relative abundance based hierarchically clustered heatmap.

**Figure 7a:**
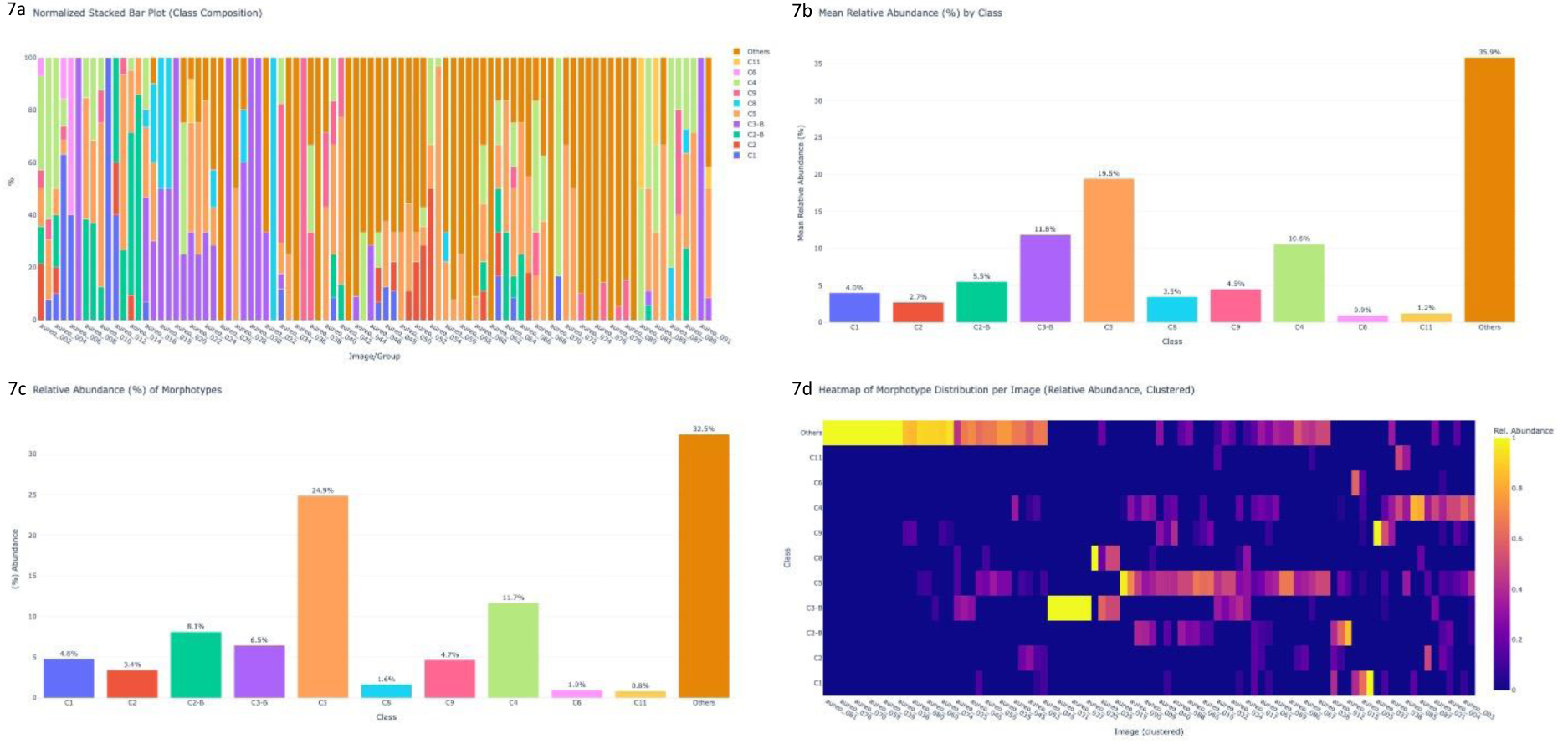
Normalized stacked bar plot of morphological composition in 90 test set images after enabling class exclusion, where C7-E, C7-F, and C7-G (filamentous morphologies) are grouped into the “Others” class. Figure 7b: Mean relative abundance (%) of morphological classes after applying class exclusion. Figure 7c: Relative abundance (%) of classes post-class exclusion. Figure 7d: Relative abundance based hierarchically clustered heatmap post-class exclusion.

#### E.2 Multi-Group Analysis Demonstrates TU_MyCo-Vision’s applicability to unseen genera

In order to assess TU_MyCo-Vision on images from different genera and test the multi-group analysis pipeline, we performed analysis on completely unseen image datasets of *C. albicans* (8), *K. phaffii* strains, *and A. niger* spores. The model could successfully detect the morphologies in the unseen data (refer to the supporting information folder “S12 data_gui_test” for the annotated images). The data analysis pipeline also successfully processed the data from all the groups and produced quantitative output and visual summaries, enabling comparative analysis among different sample groups (Figure 8). This provided an initial indication of the tool’s potential utility for morphotype quantification across taxonomically diverse fungal samples, although broader validation and detailed biological interpretations are beyond the current study’s scope.

**Figure 8a:**
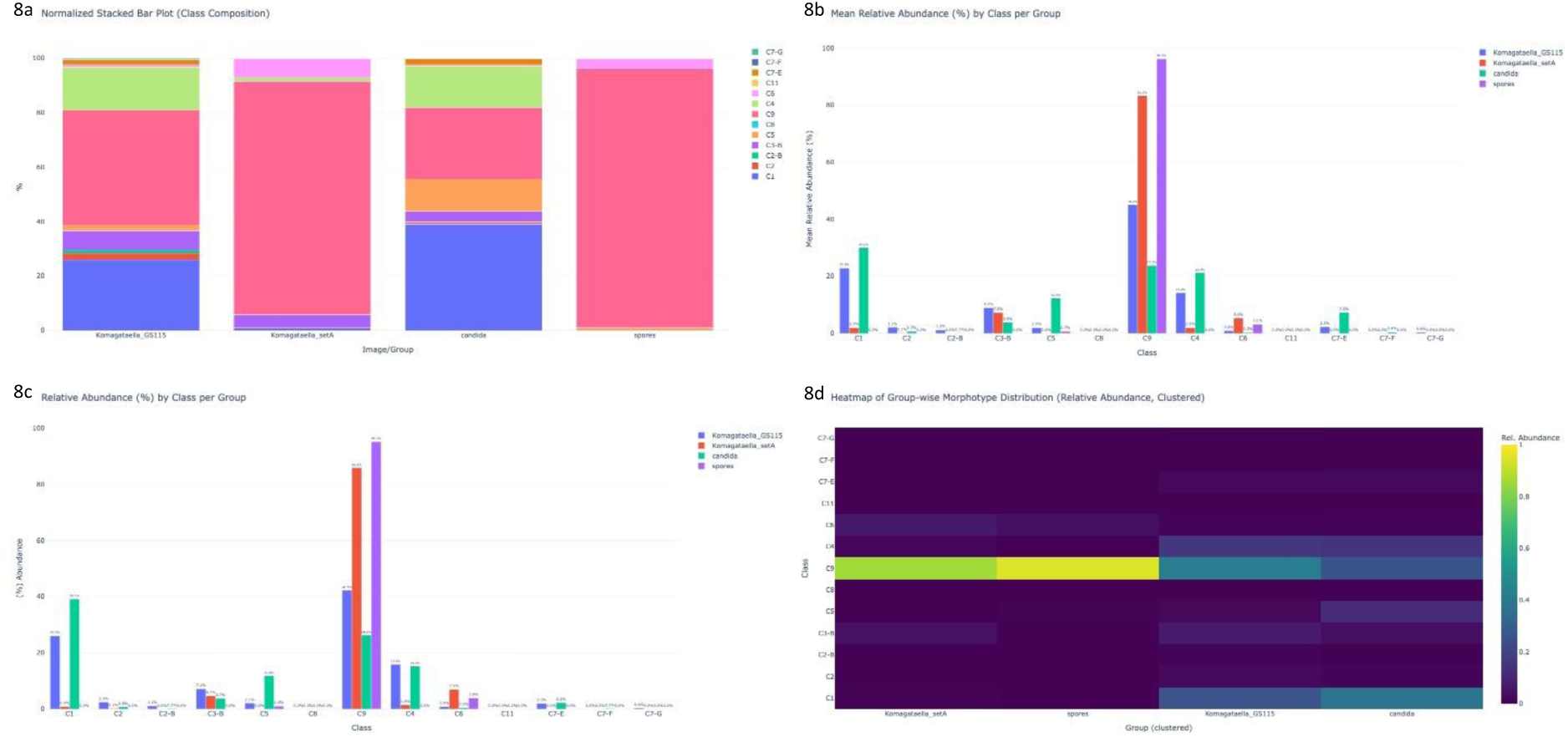
Normalized stacked bar plot representing morphological composition across four groups: Komagataella_GS115, Komagataella strain set_A, Candida, and spores. Each bar represents relative proportions of thirteen morphotypes, revealing population structure between groups. Figure 8b: Mean relative abundance (%) of each morphological class per group, providing an overview of morphological composition across groups. Figure 8c: Relative abundance (%) of morphological class within groups. Figure 8d: Relative abundance based hierarchically clustered heatmap by group.

## Discussion

In this study, TU_MyCo-Vision, an Ultralytics YOLOv11m-based object detection tool, was developed for the identification and classification of thirteen distinct fungal cell morphologies in brightfield microscopic images. The tool was designed to support investigations into the morphological plasticity of *A. pullulans*, which is a polyextremotolerant black yeast-like fungus of emerging biotechnological importance. Based on phenotypic observations using microscopic images, we defined thirteen distinct morphological forms, ranging from yeast-like to filamentous forms, and a set of special morphologies that served as the class definitions for the TU_MyCo-Vision model training. Although initially derived from *A. pullulans*, these morphologies were transferable across the genera tested and imaging contexts. We incorporated images of *A. melanogenum* and *T. reesei* into the training and validation sets to enhance the model’s generalizability.

Compared to currently available tools, such as Candescence, which focuses exclusively on *C. albicans* (8) or CCHA-YOLO, which targets clamp connections in edible fungi (9), TU_MyCo-Vision is species agnostic. The class definitions are phenotype-centric rather than species-specific or cultivation condition-specific, enhancing the tools’ adaptability to a wide range of research questions. The potential applicability of TU_MyCo-Vision to other genera and species was tested by the successful detection of morphotypes in *C. albicans* images from the Candescence dataset, *K. phaffii strains*, and on spores of *A. niger*.

Through an iterative dataset and model improvement process, we could consistently improve the performance of subsequent model families. Zulu, being the latest model family trained on a hybrid dataset of 1504 images, yielded the best-performing model. The model trained on the Zulu_split3 dataset achieved the highest performance with an average precision of 73.4%, a recall of 66.5%, and mAP@50-95 of 0.545. This indicates robust performance of the tool across all thirteen defined morphologies (refer to Table 1). Upon closer look at the validation metrics of the Zulu_split3 model, it detected cells of classes C8, C2-B, C2, C4, C3-B, and C1 with precision higher than the average precision of all classes, thus indicating superior performance for the majority of single cell classes. The model struggled with recall, i.e., detecting all the objects present in the image, especially for the filamentous classes, which could be attributed to their very complex and highly variable morphologies. This was also observed for special classes like C4 (budding events), C6 (cell aggregates), and C11 (irregular cell types), which could be the result of less training data, especially for class C11. But as the model was developed to detect the presence of different cell types, the overall performance characteristics of TU_MyCo-Vision make it a suitable tool for this purpose.

Notably, the classification of fungal cells into yeast-like, budding, and variation in hyphal morphology, such as septate and non-septate, pseudo-hyphae, is a sought-after requirement from the technician during clinical microscopic examination. It is also extremely beneficial, especially in clinical settings, to detect the presence of any fungal structure in a normally sterile bodily region, as it can be treated as proof of infection (5).

Despite overall strong performance, variability in class-wise results was observed across all model families, including Zulu. These variations stem from diverse morphologies, overlapping geometric features, filamentous classes with complex morphologies, and class imbalance within the dataset, all of which contributed to inconsistent model performance. Such challenges are common in machine learning models, where class imbalance, feature overlap, and limited data can degrade model accuracy (39). To address these issues, we employed image augmentation, which is an effective strategy to mitigate class imbalance, and adopted a smaller model architecture (YOLOv11x to YOLOv11m) to reduce overfitting on limited data (38).

To complement the core detection capabilities of TU_MyCo-Vision, we developed a cross-platform graphical user interface (GUI) to facilitate detection and downstream data analysis for users without computational skills. The GUI integrates the detection with the Zulu_s3 model and a custom data analysis pipeline that supports analysis of samples within a single group or multi-group analysis, along with additional features like class exclusion. The GUI also enables qualitative visual assessment via an integrated image viewer, which displays the original and annotated image pairs side-by-side.

We validated the GUI and data analysis pipeline with 90 images from the test dataset for single-group analysis and the class exclusion feature. For multi-group analysis testing, image sets of *C. albicans*, *K. phaffii strain GS115, and strain set_A*, and *A. niger* spores were used. This testing demonstrated the tool’s species-agnostic potential and flexibility across imaging conditions. While the primary aim of the testing was to validate the core functionality and user accessibility of TU_MyCo-Vision, we did not perform benchmarking across a wide range of fungal species or experimental conditions, as it is out of the scope of this article. Our focus was to demonstrate the potential capabilities of the tool to detect fungal cells of genera that are not part of the training dataset, highlighting its potential applicability beyond its initial training dataset.

In conclusion, TU_MyCo-Vision presents a practical, high-throughput, and user-friendly framework for automated detection and quantification of thirteen different cell types in brightfield fungal microscopic images. By integrating a SOTA (State of the Art) Ultralytics YOLOv11m-based object detector with an open-source, accessible graphical user interface and data analysis suite, this tool lowers the technical barriers for advanced functional phenotyping tools and supports flexible integration with different experimental designs. While additional optimization and systematic benchmarking across diverse fungal taxa and imaging modalities remain necessary, TU_MyCo-Vision provides a scalable and foundational solution for microscopic phenotypic analysis in fungi. Future work should focus on improving model performance, generalizability, and benchmarking across various taxa and imaging modalities.

## Acknowledgements

The computational results presented have been achieved [in part] using the Vienna Scientific Cluster (VSC). The authors gratefully acknowledge Dr. Florian Kleber and the Computer Vision Lab, Institute of Visual Computing & Human-Centered Technology, Faculty of Informatics, TU Wien, for their valuable guidance and constructive input during the model training and development. Parts of the study were supported by the TU Wien doctoral colleges, ENROL - Engineering for Life Sciences and CO_2_ Refinery.

## Data availability

All codes used during TU_MyCo-Vision development, standalone executables for Windows OS and MacOS, along with the model weight and supporting files, are available on our GitHub repository at https://github.com/cderntl/TU_MycoVision. The X-ray, Yankee, Zulu, global test dataset, and test set for GUI testing are available on our Zenodo repository at https://doi.org/10.5281/zenodo.16274901. Supporting information folders S7-S13 are available on our Zenodo repository at https://doi.org/10.5281/zenodo.16318684.

## Funding

This research was funded in whole or in part by the Austrian Science Fund (FWF) [10.55776/P 35642]. For open access purposes, the authors have applied a CC BY public copyright license. Author Johannes. T. Zwerus gratefully acknowledges financial support from the Austrian Federal Ministry of Economy, Energy and Tourism, the National Foundation for Research, Technology and Development, and the Christian Doppler Research Association.

## Author contribution

KJD Conceptualization, Data Curation, Formal Analysis, Investigation, Methodology, Software, Visualization, Writing – Original Draft Preparation

MS Methodology, Software

CD Investigation

ZAQ Investigation

JTZ Investigation

JK Investigation

MH Investigation

RS Investigation

AMA Resources

RLM Resources, Supervision

CZ Funding Acquisition, Supervision, Writing – Review & Editing

## Supporting Information

Supporting information folders S7-S13 are available on Zenodo repository at https://doi.org/10.5281/zenodo.16318684”.

S1 Table 1. Detailed medium composition.

S2 Table 2. Distribution of objects across all classes in training and validation sets, in all five splits, and three dataset families.

S3 Table 3. Class-wise object distribution in the global test dataset.

S4 Table 4. Hyperparameter set post-exploratory search.

S5 Table 5. Tuned Hyperparameters.

S6 Table 6. Hyperparameters (Zulu model family).

S7 Folder V1. Validation metrics and visualizations (model V1).

S8 Folder hyperparameter_tuning. Results of hyperparameter tuning.

S9 Folder xray. Validation metrics and visualizations (model X-ray).

S10 Folder zulu. Validation metrics and visualizations (model Zulu).

S11 Folder yankee. Validation metrics and visualizations (model Yankee).

S12 Folder data_gui_test. Predictions, single-group and multi-group analysis data.

S13 Folder validation_global_set. Performance visualization of all models validated on the global test dataset.

S14 Table 7. validation_metrics_global_test_set. Detailed validation metrics of all models validated on global test dataset.

## Notes

https://github.com/cderntl/TU_MycoVision

https://doi.org/10.5281/zenodo.16274901

https://doi.org/10.5281/zenodo.16318684

### Competing Interest Statement

The authors have declared no competing interest.

